# Redox-constrained microbial ecology dictates nitrogen loss versus retention

**DOI:** 10.1101/2025.07.14.664727

**Authors:** Jemma Fadum, Xin Sun, Emily Zakem

## Abstract

Microorganisms drive biogeochemical cycling. Therefore, examining environmental change through the lens of microbial ecology is particularly useful for developing a mechanistic understanding of the biogeochemical consequences and feedbacks of perturbations to ecosystems. When aquatic systems with deep anoxic waters undergo eutrophication, the resulting surface productivity impacts the anaerobic microbial community below. The increase in sinking organic carbon can shift the anaerobic community function from inorganic nitrogen (N) loss to N retention, amplifying eutrophication as a positive feedback. However, we lack a mechanistic understanding of this transition, which is critical for anticipating these impacts in aquatic environments where microbial community composition is unknown. Here, we provide a first-principles, quantitative model of this transition from N loss to retention by linking ecological dynamics to the energetics underlying microbial metabolisms. We develop and analyze an ecosystem model in which redox chemistry constrains the traits of key anaerobic N-cycling microbial functional types: denitrification, dissimilatory nitrate reduction to ammonium (DNRA), and anaerobic ammonium oxidation (anammox). The model captures the transition from N loss to N retention with increasing organic carbon supply, consistent with previous observations for specific systems and species. Results identify characteristics of the microbial community composition at the ‘net zero N loss’ point at which N loss balances N retention, providing testable hypotheses for sequencing data and other observations. By tying microbial ecological dynamics to environmental chemical potential, results provide a broadly applicable framework for improving predictions of the biogeochemical impacts of eutrophication alongside deoxygenation and other ecosystem perturbations.

## INTRODUCTION

Aquatic ecosystems face a myriad of impacts related to global change, from anthropogenically driven changing climate patterns to more localized disturbances. Such impacts, whether local or global, often manifest through changes in microbial community function. One ubiquitous impact on both freshwater and coastal ecosystems is eutrophication, characterized broadly by increased algal productivity and accompanying ecosystem changes caused by excessive nutrient supplies. Excess nutrients can come from innumerable point and nonpoint sources such as the overapplication of fertilizers, untreated (or undertreated) municipal waste, and net pen aquaculture, among other sources. Nutrients may take the form of inorganic compounds (e.g., nitrate, ammonium, and phosphate), as well as more complex and diverse organic compounds, which we consider collectively here as organic matter (OM). OM loading is often a key driver of eutrophication (Deininger and Frigstad 2019) because oxidation of the carbon (C) in OM by heterotrophic metabolisms often releases bioavailable inorganic nutrients that fuel primary production.

The algal blooms that develop in eutrophic systems can subsequently produce hypoxic (low oxygen) or anoxic (often undetectable oxygen) zones in the water column as decomposing algal biomass and other OM sinks below the ventilated surface waters. In these deeper waters, oxygen supply can be insufficient to meet the demands of aerobic metabolism (Diaz 2001). Often called “dead zones” for their inability to support marine megafauna, these low oxygen environments are still teeming with microbial life (Wright, Konwar, and Hallam 2012). When oxygen is unavailable, microbial communities operate anaerobically by using oxidized forms of inorganic nitrogen (N) and other molecules as electron acceptors.

In anoxic waters, the combination of active N-cycling microbial metabolisms determines whether bioavailable N is lost or retained in the ecosystem. Both aerobic and anaerobic metabolisms can be biogeochemically relevant when oxygen is supplied at low levels (Garcia-Robledo et al. 2017, Canfield and Kraft 2022). When oxygen supply is negligible, three anaerobic metabolisms (or else a subset of the three) can be expected to dominate anoxic water column N-cycling: denitrification (NO_3_^-^ → NO_2_^-^ → NO → N_2_O → N_2_), anaerobic ammonium oxidation (anammox; NH_4_^+^ + NO_2_^-^ → N_2_), and dissimilatory nitrate reduction to ammonium (DNRA; NO_3_^-^ → NO_2_^-^ → NH_4_^+^). Denitrification converts bioavailable species of N to gaseous forms, producing both dinitrogen gas (N_2_) and nitrous oxide (N_2_O), a potent greenhouse gas with a warming potential 300 times that of CO_2_ (IPCC 2013). Anammox also results in the loss of bioavailable N but does not produce N_2_O. In contrast, DNRA uses nitrate (NO_3_^-^) or nitrite (NO_2_^-^) as an electron acceptor to produce ammonium (NH_4_^+^), thus retaining bioavailable N in the ecosystem, though also producing negligible amounts of N_2_O (Bleakley and Tiedje 1982, Rütting et al. 2011)

The degree to which bioavailable N is lost or retained in a strictly anoxic aquatic ecosystem, where aerobic metabolisms are negligible, is thus primarily determined by the proportion of active denitrification, anammox, and DNRA. Denitrification and DNRA utilize similar types of reduced substrates as electron donors, such as OM if operating heterotrophically. Therefore, the populations carrying out these metabolisms can be considered to be competing for both NO_x_ and OM. In contrast, chemoautotrophic anammox is often understood to exist in syntrophy with denitrification, relying on the NH_4_^+^ excreted from the remineralization of OM by denitrifying heterotrophs (Koeve and Kähler 2010, Babbin et al. 2014), though it has also been observed to co-occur with DNRA (Lam et al. 2009). Despite competition for often-limited resources, the coexistence of DNRA, denitrification, and anammox has been observed in both marine and freshwater environments (e.g., Lam et al. 2009, Kalvelage et al. 2013, De Brabandere et al. 2014, Roland et al. 2018, among others).

Previous research has demonstrated that, generally, DNRA is more favorable when microbial growth is NO_x_-limited whereas denitrification is more favorable when microbial growth is OM-limited (Cole and Brown 1980, Tiedje et al. 1983, Strohm et al. 2007, Algar and Vallino 2014, van den Berg et al. 2016, Jia, Winkler, and Volcke 2020), though exceptions to this pattern can occur in metabolically flexible organisms (Vuono et al. 2019). A transition to the dominance of DNRA (i.e., the N retention pathway) in increasingly OM-rich environments provides a positive feedback to eutrophication (Fig. 1). As eutrophication expands, elevated OM sinking from increasingly productive surface waters may transition the ecosystem from denitrification dominant to DNRA dominant, thus retaining bioavailable N and further spurring eutrophication by supplying inorganic N which supports primary productivity. In the other direction, if supply of OM decreases, the system may shift towards a more denitrification dominant state, increasing bioavailable N loss, thus potentially reducing surface primary production and subsequently the supply of OM to anoxic zones below. This feedback is particularly important in aquatic ecosystems where primary productivity is N-limited (for example, roughly half of the surface ocean and some inland bodies of water, Moore et al. 2013, Fadum and Hall 2023).

**Figure 1.**
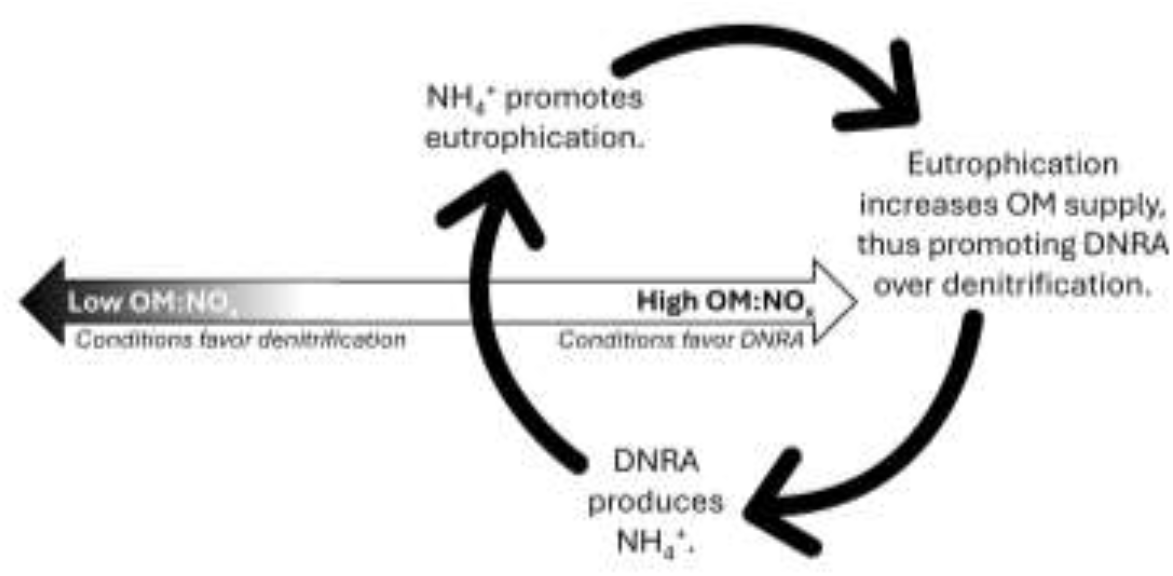
Conceptual diagram of the positive feedback where DNRA is promoted by eutrophication while simultaneously supplying nutrients which fuel eutrophication.

In order to make accurate projections of N loss in changing anoxic environments, a mechanistic, quantitative understanding of the controls on these key metabolisms is required. Jia et al. (2020) provided such a quantitative understanding using Resource Competition Theory to interpret observations (Tilman 1982). They demonstrated that measured denitrification and DNRA outcomes can be predicted by the ratio of organic C to NO_3_^-^ supply and also identified the stable coexistence of denitrification and DNRA. However, the model parameters were set by measured traits of specific microbial populations. Given the taxonomic breath of both denitrification and DNRA-performing organisms (Kraft, Strous, and Tegetmeyer 2011), a more general model is needed in order to predict changes in unobserved, unculturable, and future-adapted communities.

One way forward is to link population traits to fundamental underlying constraints, foregoing species-specific descriptions in favor of more general and broadly applicable trait-based descriptions of functional types (Litchman et al. 2007, Kiørboe, Visser, and Andersen 2018). For metabolically diverse prokaryotes, the key redox reaction fueling a metabolism provides one such fundamental constraint (Vallino, Hopkinson, and Hobbie 1996, Rittman and McCarty 2001, Zakem, Polz, and Follows 2020). In line with this approach, Algar and Vallino (2014) optimized for maximum entropy production (MEP) among N-cycling metabolisms in a coastal sediment environment, which, like Jia et al. (2020), explained the niche differentiation of denitrification and DNRA according to the ratio of organic C to NO_3_^-^ supply. However, the study focused on sediments, rather than the water column, where the supply mechanisms and magnitudes of essential substrates differ. In addition to the fundamental differences between sediments and the water column environment, a model framework devised for the water column could allow for integration with multi-dimensional circulation models of aquatic environments, including the biogeochemical models used for climate change projections.

Here we link environmental chemical potential (i.e., nutrient availability) to the ecological dynamics of populations carrying out the anaerobic N-cycling metabolisms that dictate the balance between inorganic N loss and retention in anoxic water columns. We develop and analyze an ecological model with microbial functional types that are parameterized with the underlying free energies and stoichiometries of the redox reactions that fuel growth (Zakem, Polz, and Follows 2020), providing a theoretically grounded quantitative framework to study microbial N-cycling, in contrast to a species-specific approach. We focus on the ecological interactions between microbial populations (functional types) carrying out the three dominant anaerobic metabolisms (denitrification, DNRA, and anammox, Fig. 2). Like Jia et al. (2020), we use Resource Competition Theory to understand the ecological outcomes. The tradeoff resulting in the niche differentiation of denitrification versus DNRA embedded in the underlying energetics emerges in the traits of the functional types, demonstrating the ability of the modeling strategy to link chemical potential to trait-based descriptions of microbial communities. We then analyze the transition between N loss and N retention from a microbial ecological perspective and identify the microbial community composition at the transition point. This provides testable hypotheses for sequencing datasets and other observations.

**Figure 2.**
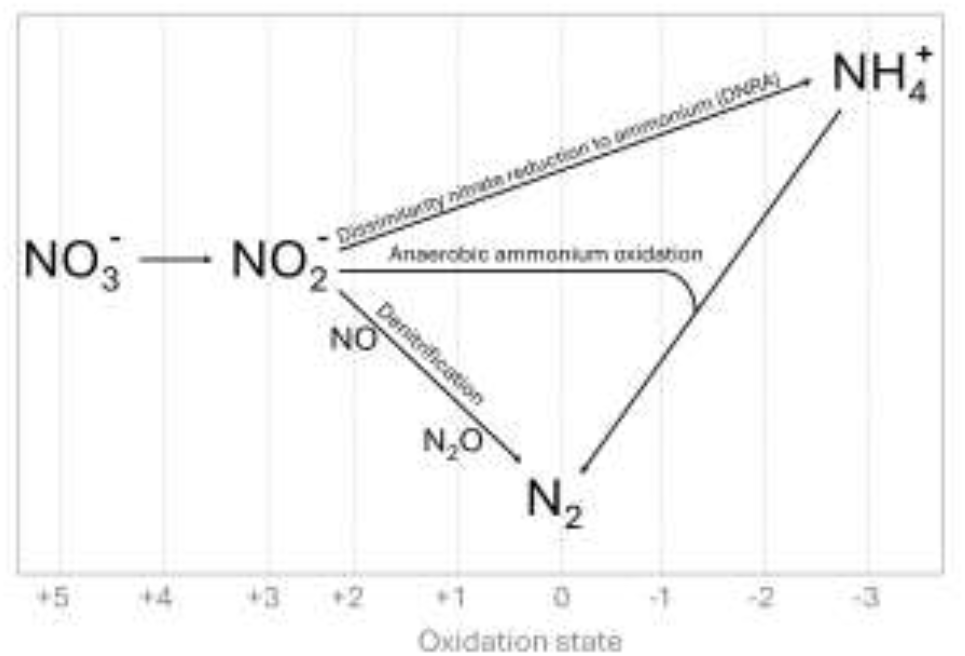
Conceptual diagram of the key anaerobic N-cycling pathways. The fate of nitrate (NO_2_^-^) is determined by three main metabolisms (denitrification, anammox, and DNRA), which are represented by microbial functional types in the ecosystem model.

## METHODS

### Redox-constrained microbial functional types

We employ an established framework that uses the underlying redox reactions fueling metabolisms to quantitatively describe the N-cycling microbial functional types responsible for the three main anaerobic N-cycling pathways: denitrification, DNRA, and anammox (Fig. 2; Zakem et al. 2020, Sun et al. 2024). Specifically, we develop and analyze a heterotrophic DNRA functional type and add it to previously developed functional types for heterotrophic denitrification and chemoautotrophic anammox (Sun et al. 2024). Briefly, substrate (OM and inorganic N) requirements for the heterotrophic functional types are quantified using the half reactions of oxidized and reduced substrates and an estimated proportion of electrons devoted to biomass synthesis versus respiration (Rittman and McCarty 2001). This reflects the free energies of the reactions with consideration of the departure from standard state due to typical concentrations of these solutes in anoxic waters. As in previous work (Sun et al. 2024), we align the magnitudes of the biomass yields for all heterotrophic functional types with a range of empirically estimated biomass yields (or, C use efficiency) on organic C for natural aquatic environments (0.2—0.3 mol biomass C per mol C; Sinsabaugh et al. 2013). Thus, the relative values of the yields reflect the underlying redox chemistry, while the absolute values reflect this empirically estimated baseline. See Sun et al. (2024) for full, detailed methodology for determining yields. See Supporting Information Text S1 for DNRA-specific details.

As the first step of DNRA and denitrification (NO_3_^-^ → NO_2_^-^) is not distinguishable from a redox perspective, we focus on the competition for NO_2_^-^, the critical intermediate that determines the fate of bioavailable N (Algar and Vallino 2014, Kraft et al. 2014, van den Berg et al. 2017, Tracey et al. 2023). Due to the known modularity of denitrification (Zhang et al. 2023), we resolve the denitrification pathway with two functional types that each carry out a single-step pathway (NO_2_^-^ → N_2_O and N_2_O → N_2_). We use a previously published parameterization for the chemoautotrophic anammox functional type that relies on measured yields (Lotti et al. 2014, Buchanan et al. 2025). We represent OM simply as one bulk pool with typical marine stoichiometry (C_6.6_H_10.9_O_2.6_N, Anderson 1995), though the methodology is sufficiently flexible to consider different stoichiometries and specific substrates.

This results in the following metabolic budgets, with each normalized to the synthesis of N-based biomass (with stoichiometry C_5_H_7_O_2_N; Rittman and McCarty 2001, Zakem et al. 2020), here in simplified form (neglecting water and proton balancing) for conciseness. For heterotrophs, the budgets reflect the baseline biomass yield range of 0.2—0.3 mol biomass C per mol organic C, here written in terms of the midpoint (0.25 mol/mol), which is consistent with the empirically estimated 0.26 mol/mol for aquatic environments (Sinsabaugh et al. 2013):

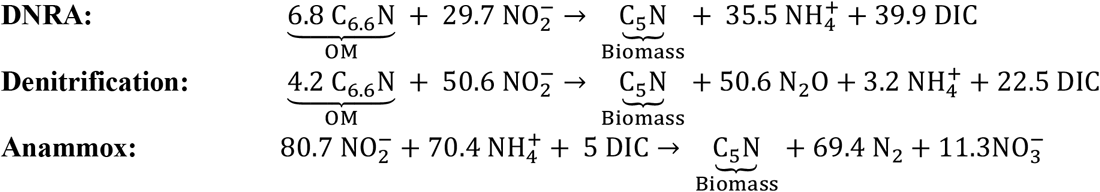

### Ecosystem model

A virtual chemostat model is seeded with the anaerobic functional types (denitrification, DNRA, and anammox) and supplied with varying incoming concentrations of OM and NO_2_^-^ (Supporting Information Table S1). Ammonium (NH_4_^+^) is also supplied to the chemostat at a rate that represents the NH_4_^+^ released by implied nitrate (NO_3_^-^) reduction (Supporting Information Text S2). We numerically integrate the model to equilibrium for each combination of incoming concentrations.

In addition to the biomass yields (Table 1), the functional type descriptions require parameters dictating substrate uptake kinetics and population loss due to mortality. As in previous work (Zakem et al. 2020, Sun et al. 2024), we have kept the substrate uptake kinetic parameters and loss rate parameters the same for all types (Supporting Information Table S2) in order to examine the competitive outcomes produced by differences in the yields, which reflect the differences in the underlying redox chemistry. Specifically, the growth rate μ (d^-1^) for each functional type *i* on required resource *j* (*R*_*j*_) is calculated using Leibig’s law of the minimum as:

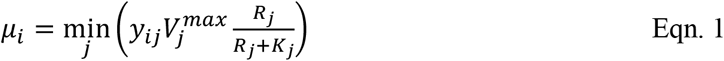

so that the maximum growth rate (*μ*_*max*_) is the product of the yield and the maximum uptake rate *V*^*max*^ (d^-1^), where *V*^*max*^ is the same for each functional type. The loss rates are set by the chemostat dilution rate. However, because of the additional uncertainty in comparing heterotrophic and chemoautotrophic metabolisms, we later forego this assumption for anammox and instead consider that it may have optimized its cellular machinery to achieve a different competitive outcome.

**Table 1.** Comparison of yields (mols biomass N per mol resource). The range of DNRA and denitrification yields and resource subsistence concentrations (R*) reflects the range in the baseline biomass yield with respect to organic N (0.2-0.3 mols of biomass per mol N in OM oxidized). Anammox *y*_*N*_ (and resulting NO_2_^-^*) from (Lotti et al. 2014).

| Metabolism | $y_{\text{OM}}$ | $y_N$ | $\text{OM}^*$ | $\text{NO}_2^*$ |
| --- | --- | --- | --- | --- |
| **DNRA** | 0.11-0.18 | 0.026-0.042 | 0.080-0.12 | 0.0032-0.0054 |
| **Denitrification ($\text{NO}_2^- \rightarrow \text{N}_2\text{O}$)** | 0.19-0.29 | 0.015-0.024 | 0.048-0.074 | 0.0058-0.0097 |
| **Anammox** | NA | 0.0133 | NA | 0.0123 |

Finally, we use Resource Competition Theory to interpret the results (Tilman 1982). In the resulting resource ratio diagrams, the consumption vectors (CVs) are the ratios of the resource yields (i.e., *y*_N_/ *y*_OM_), representing the ratio of OM to NO_2_^-^ consumption for the heterotrophs.

## RESULTS

### DNRA-denitrification tradeoff reflects redox chemistry

In a steady-state ecosystem, where microbial growth is faster than changes in the environment, the competitive ability of each functional type for a nutrient is set by its resource subsistence concentration (R*), which is the minimum concentration that can sustain a population in the environment, and the concentration to which the population will deplete that resource when its growth is limited by that resource (Tilman 1982). In our model, each subsistence concentration is set by the population’s specific loss rate (i.e., the chemostat dilution rate), uptake kinetics (here, the maximum uptake rate and the half saturation constant, Supporting Information Table S2), and the biomass yield (Zakem et al. 2020, Sun et al. 2024; Table 1). For the competing heterotrophs, the subsistence concentrations for NO_2_^-^ (NO_2_*) and OM (OM*) differ solely because of the differences in the redox-informed biomass yields (Table 1). The biomass yields are calculated from the metabolic budgets (Zakem et al. 2020; Sun et al. 2024, Supporting Information Text S1) and are inversely related to the subsistence concentrations (Table 1). A larger yield/smaller R* for a nutrient means a given functional type is more competitive when that nutrient is limiting.

Results show that, with respect to the consumption of NO_2_ ^-^, the denitrifying functional type has a greater OM yield (*y*_OM_) than DNRA (Table 1), which is consistent with empirical observations of denitrification dominance under OM limiting conditions. Conversely, DNRA has a greater NO_2_^-^ yield (*y*_N_) than denitrification, which is consistent with DNRA’s observed dominance under NO_x_ limited conditions. Anammox has the lowest NO_2_^-^ yield (∼18% lower than denitrification), reflecting the slow growth and high energetic cost of chemoautotrophy (Mccollom and Amend 2005).

These yield estimates demonstrate that the metabolic tradeoff between OM and N utilization embedded in the redox reactions of denitrification and DNRA emerges in the theoretically modeled traits, without a dependency on other physiological or ecological factors. This is consistent with our hypothesis that fundamental constraints underlying metabolisms (i.e., chemical potential energy) predominantly govern the fitness of populations carrying out metabolisms that are sufficiently comparable (i.e., heterotrophic denitrification versus DNRA, considering that both gain energy from the oxidation of the same OM). Differences in kinetic parameters, such as a higher affinity for NO_2_^-^ for DNRA, are likely to evolve in accordance with these metabolic niches (as reflected in the measured traits in Jia et al. 2020), but we hypothesize that traits such as uptake affinity reflect a cellular-level optimization and thus are unlikely to ‘override’ the fundamental redox-level constraints for comparable metabolisms (i.e., heterotrophs). We next examine the outcomes of the interactions of these functional types in the ecosystem model.

### Ecological outcomes

We incorporate the denitrification, DNRA, and anammox functional types into a simplified ecosystem model in order to: 1) illustrate how the redox-based tradeoff between higher competitive ability for OM versus NO_2_^-^, for denitrification and DNRA, respectively, shapes community composition across variable resource availability, 2) examine the metabolic niche for anammox, and 3) examine how the ecological outcomes relate to the biogeochemical outcome: the balance between N retention versus N loss.

Resulting steady-state model simulations show the variation in the ratio of active denitrification to DNRA as a function of OM:NO_2_^-^ supply rates (Fig. 3). To more clearly and quantitatively compare the biomasses, rates, and geochemistry, a subset of these simulations is illustrated in one dimension (a single NO_2_^-^ supply rate with varied OM supply rates) (Fig. 4). In low OM:NO_2_^-^ space, denitrification competitively excludes DNRA, and in high OM:NO_2_^-^ space, DNRA competitively excludes denitrification. These zones of exclusion are set by consumption vectors (CV), which represent the relative consumption rates of OM versus N by the functional types. DNRA competitively excludes denitrification at OM:NO_2_^-^ supply rates above its CV, which is equal to the ratio of its biomass yields for OM and N (Fig. 3A-B). In contrast, denitrification competitively excludes DNRA at OM:NO_2_^-^ supply rates below its CV. Stable coexistence of denitrification and DNRA, despite their competition for both substrates, occurs where the two functional types are co-limited by both OM and NO_2_^-^, between the two CVs (Fig. 3A-B, Fig. 4A).

**Figure 3.**
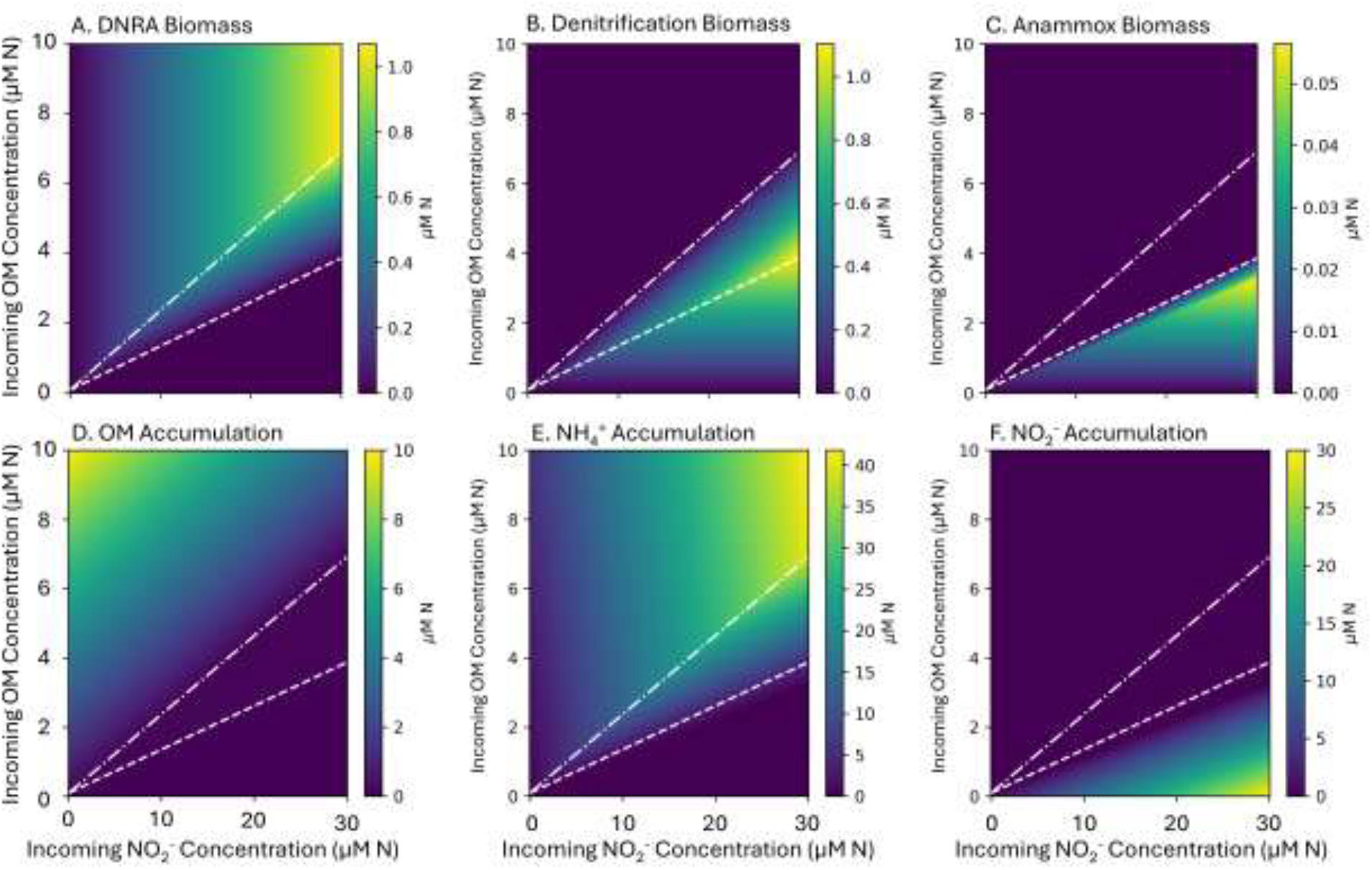
Multiple steady-state solutions of the ecosystem model with anaerobic N-cycling metabolic functional types across varying supply of OM and NO_2_^-^, showing biomass of A) DNRA, B) Denitrification (NO_2_^-^ → N_2_O + N_2_O → N_2_) and C) Anammox. Chemostat accumulation (steady-state concentrations) of D) OM, E) NH_4_^+^ with F) NO_2_^-^. The white lines indicate the consumption vectors for DNRA (− • − • −) and denitrification (−--).

**Figure 4.**
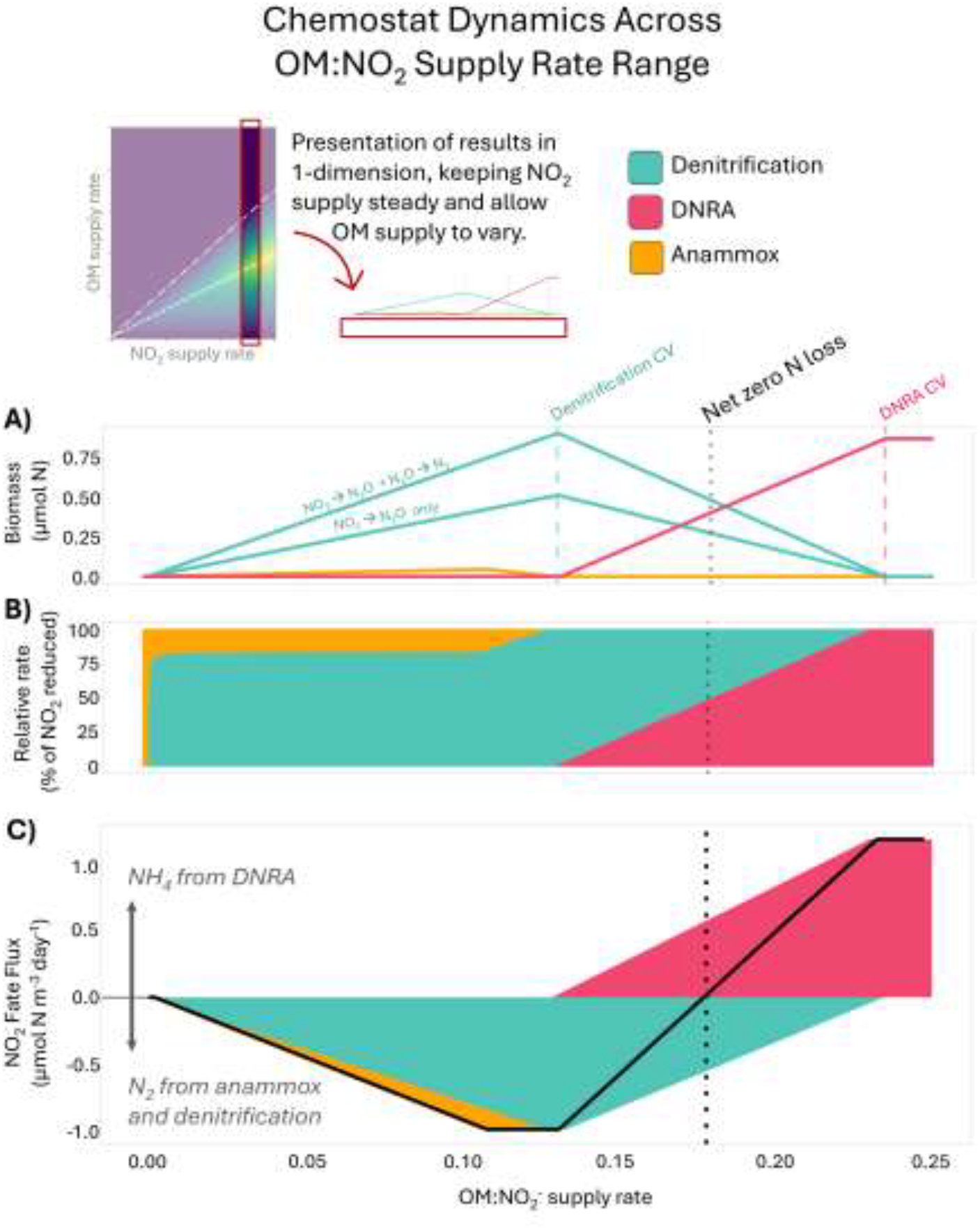
Ecological and biogeochemical outcomes governing the fate of NO_2_^-^ in the ecosystem model as a function of the relative supply of OM to NO_2_^-^ (here specifically with incoming NO_2_^-^ concentration of 25 μM N and variable OM supply rate). A) Biomasses of DNRA, denitrification, and anammox functional types, B) Percentage of NO_2_^-^ reduced via the three functional types, C) N_2_ and NH_4_^+^ production, identifying point of net zero N loss.

These results are consistent with those of Jia et al. (2020) yet without reliance on species-specific, measured parameters. Thus, the results of Jia et al. (2020) serve as a test of the ability of our framework to successfully connect fundamental constraints (underlying energetics) to observed microbial communities, thus supporting the utility of this framework for anticipating dynamics in unobserved microbial communities where the species composition is unknown.

These competitive outcomes align with the patterns of nutrient accumulation. We consider a nutrient ‘accumulating’ when its concentration is above the subsistence concentration of any microbial functional type. In low OM:NO_2_^-^ space, NO_2_^-^ (the electron acceptor) accumulates due to insufficient OM (the electron donor) to make use of the entire electron acceptor pool, Conversely, in high OM:NO_2_^-^ space, OM accumulates due to insufficient NO_2_^-^ to make use of the entire electron donor pool. In the coexistence regime, neither OM nor NO_2_^-^ accumulates because both limit microbial growth (Fig 3D, Fig. 3F).

### The role of anammox

In the default version of the model, anammox is less competitive for NO_2_^-^ than the heterotrophs because of its very low NO_2_^-^ yield (*y*_*N*_) (Table 1). The resulting anammox biomass is generally an order of magnitude lower than both DNRA and denitrification biomass (Fig. 3A-C, Fig. 4A), consistent with the observed low growth rate and low biomass accumulation in some natural environments (Kartal et al. 2012). Anammox is only sustained in the OM-limited regime at low OM:NO_2_^-^ supply rates, where there is sufficient NO_2_^-^ as well as sufficient NH_4_^+^ to serve as the electron donor. In this regime, anammox and denitrification maintain stable coexistence due to syntrophy, with the heterotrophs supplying NH_4_^+^ from the remineralization of the OM (Fig. 3B-C, Fig. 4A). In this regime, anammox is responsible for ∼20% of NO_2_^-^ uptake (Fig. 4B) in the chemostat and ∼26% of N_2_ production (Supporting Information Fig. S1), which is consistent with the established geochemical calculations of the contribution of anammox to N loss in the ocean (∼28%) reflecting control by the OM stoichiometry (Koeve and Kähler 2010; Babbin et al. 2014). At higher OM:NO_2_^-^ supply, once NO_2_^-^ becomes depleted, anammox transitions from a state of NH_4_^+^ limitation to NO_2_^-^ limitation. Because the *y*_N_ of anammox is too low (and thus NO_2_^-^* is too high) to compete with DNRA for NO_2_^-^, the chemoautotroph is eventually excluded as NO_2_^-^ becomes limiting.

However, given some observations of anammox co-occurring with DNRA in the ocean (Kalvelage et al. 2013), we also considered the possibility that anammox may be a better competitor for NO_2_^-^ than the heterotrophs. Though we hypothesize from an evolutionary perspective that redox constraints should inform ecological niches of comparable metabolisms, such as heterotrophic denitrification and DNRA fueled by the same electron donor, our framework involves more uncertainty for comparing the niches of heterotrophs versus chemoautotrophs. Despite the low *y*_N_ of anammox in the default model, which reflects the high energetic cost of chemoautotrophy, it is plausible that anammox is able to devote more of its cellular machinery to NO_2_^-^ acquisition than the heterotrophs because it does not need to process complex organic substrates. If this cellular-level optimization is sufficient for anammox to have a lower NO_2_^-^* than DNRA and denitrification (Supporting Information Fig. S2-S4), then the anammox functional type would be a superior competitor for NO_2_^-^ (despite observations and previous work suggesting otherwise, Lotti et al. 2014; van den Berg et al. 2016). We conduct this model experiment and find that if this is the case, anammox can access NO_2_^-^ throughout the model domain and coexist with DNRA, with DNRA supplying NH_4_^+^ throughout the N-limited regime (Supporting Information Fig. S2-S3). In this scenario (NO_2_*_*anammox*_ < NO_2_*_*DNRA*_< NO_2_*_*denitrification*_), the model produces a domain of stable coexistence of all three functional types (Supporting Information Fig. S2-S3). This simulates the observed co-occurrence of all three functional types (Kalvelage et al 2013), although other mechanisms could also allow for this coexistence (see discussion). Overall, this analysis generates a targeted question for observations and experiments about the competition between anammox and heterotrophs for NO_2_^-^.

### The balance between N loss and N retention

The outcome of the ecological interactions of the anaerobic N-cycling functional types results in varied fate of NO_2_^-^ across the resource ratio space. In high OM:NO_2_^-^ space, NH_4_^+^ production from DNRA dominates, and N is retained in the system (Fig. 4C). In low OM:NO_2_^-^ space, N_2_ production from denitrification and anammox dominates, and N is lost from the system. Critically, the coexistence regime spans a range of NO_2_^-^ fate, from N loss to N retention. Thus, the model indicates that the presence of active denitrification does not necessarily mean that there is more N loss than N retention, and the presence of active DNRA does not mean that there is more N retention than N loss. Next, we examine the resource ratio that allows for an exact balance between N retention and N loss, for anaerobic metabolisms alone, and how this relates to the anaerobic microbial community structure.

### Net zero N loss

We identify the resource ratio (OM:NO_2_^-^) where NH_4_^+^ produced from DNRA (i.e., retention of inorganic N) balances N_2_ production from denitrification and anammox (i.e., loss of inorganic N), which we refer to as ‘net zero N loss’ (Supporting Information Text S3). This balance occurs within the DNRA and denitrification coexistence regime, where anammox is excluded, and where DNRA and denitrification are equally competitive for their shared electron acceptor, thus able to reduce equal amounts of NO_2_^-^ (Fig. 4B). In the model equations, this balance is:

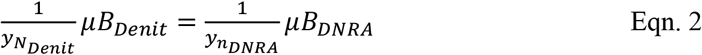

If we assume that both populations have similar steady-state growth rates, μ, as they do in the chemostat, which is likely the case in the environment if they are relatively similarly sized cells and thus subject to similar grazing rates, we can then solve for the OM:NO_2_^-^ supply ratio that results in net zero N loss as a function of only the biomass yields for the two populations as:

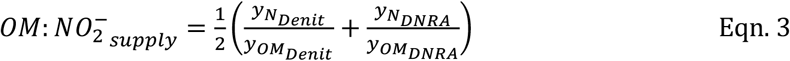

Because the ratios of the yields set the consumption vectors (CVs), net zero N loss coincides with the geometric mean of the two CVs (Fig. 4). Along this mean, the biomass of the population carrying out DNRA is only slightly less than the total biomass of the two combined populations carrying out denitrification (NO_2_^-^ → N_2_O and N_2_O → N_2_). Thus, the model suggests that nearly equal amounts of DNRA-associated and denitrification-associated biomass in the environment may indicate net (or near net) zero N loss, for a strictly anoxic environment. With our range of baseline biomass yields (Table 1), this net zero N loss point is between 0.1729 - 0.1820 mol N (in OM) per mol NO_x_ supplied. Because the C in the OM is what ultimately limits the heterotrophs, this range would vary quantitatively but not qualitatively with the C:N stoichiometry of the OM supplied.

## DISCUSSION

We have provided a broadly applicable framework for understanding the ecological dynamics of anaerobic N-cycling microbial communities in anoxic water columns, both freshwater and marine. Specifically, we demonstrate that by considering the free energy of the underlying half reactions, we are able to simulate the ecological outcomes of the interactions between populations carrying out DNRA versus denitrification that have been theoretically hypothesized and observed and modeled for specific taxa (Tiedje et al. 1983, Algar and Vallino 2014, Jia, Winkler, and Volcke 2020). Our model results are also consistent with the results for the sediment environment of Algar and Vallino (2014), although we use an alternative modeling framework and methodology that relies on the outcome of modeled ecological interactions rather than optimization. Therefore, our model is amenable to incorporation into more complex, multidimensional biogeochemical models that typically already include explicit microbial populations (e.g., phytoplankton and zooplankton). If incorporated into multi-dimensional global biogeochemical ocean models, which do not consider DNRA (and perhaps more significantly, its competition with denitrification), the presented framework may provide additional insights into global N-cycling. The model may also offer key insights into both contemporary and future ecosystem function where environmental parameters and community composition data are unavailable (i.e., much of the global ocean and inland waters in historically and chronically understudied regions; Anil et al. 2018, Smits et al. 2025).

Our model links mechanistic microbial ecological dynamics to geochemical quantities that are frequently measured in both freshwater and marine environments. For example, the speciation of accumulated inorganic N is reflective of the dominant functional type (Fig 3E-F, Supporting Information Fig. S5). This suggests broad signatures of microbial metabolism in the accumulation patterns of inorganic N species: denitrification and anammox when NO_x_ accumulates, DNRA when NH_4_^+^ accumulates, and coexistence when neither accumulates. These same patterns have been observed in natural ecosystems such as in oligotrophic OMZs (where OM:NO_2_^-^ supply rates are putatively low, mirroring the left side of the panels of Fig. 4) and the hypolimnion of eutrophic lakes (where OM:NO_2_^-^ supply rate would be expected to be high, mirroring the right side of the panels of Fig. 4). In oligotrophic OMZs, NO_2_^-^ accumulation concomitant with denitrification and/or anammox dominance has been observed (Ward et al. 2009, Dalsgaard et a. 2012, Kalvelage et al. 2013, among others) whereas NH_4_^+^ accumulation and DNRA dominance has been observed in the hypolimnion of stratified lakes (Pajares et al. 2017, Fadum et al. 2024). Therefore, our model captures broadscale, observed patterns from a first-principles understanding of microbial community function.

Because functional microbial biomass is resolved explicitly, results provide additional hypotheses that can be tested with more direct observations of microbial communities. For example, the model provides testable hypotheses that could generate new insights into the amassed wealth of sequencing data from diverse aquatic environments. The model mechanistically links microbial ecology to biogeochemical fluxes by relating the relative concentrations of functional type biomasses to nutrient transformation rates. Specifically, the modeled biomasses can be compared to gene-based estimates of abundances by converting functional biomass (the concentration of C- or N-based biomass in the water) to abundance using an estimate of the cell quota (amount of C or N per cell) and an estimate of the number of gene copies per cell. The bulk N transformation rates in the model (e.g., denitrification) can also be converted to a per cell rate using these conversion factors. Thus, microbially explicit biogeochemical modeling brings the aquatic sciences closer to being able to use gene expression to estimate both rate processes and elemental fluxes (Smets et al. 2016, Kleiner et al. 2017). Though this quantitative alignment involves uncertainty (Meiler et al. 2022), the model may also provide qualitative descriptions of dominant pathways and therefore add important context to genomic datasets where genes multiple NO_x_ reducing pathways are present.

A specific question raised by our analysis that could be tested with observational methods is in regard to the outcome of the competition between DNRA and anammox for NO_2_^-^. Anammox is competitively excluded in the NO_2_^-^ limited regime by DNRA in the default model (Fig. 3, 4). This suggests that anammox is only sustainable when NO_2_^-^ does not limit microbial growth, which we speculate is the most likely outcome. However, when anammox was given a lower NO_2_* (and thus higher *y*_N_), it was in principle able to outcompete DNRA (Supporting Information Fig. S2-S3), yet its requirement for NH_4_^+^ meant that it relied on NH_4_^+^ supplied from DNRA, and thus anammox coexisted with DNRA in the NO_2_^-^ limited regime. Therefore, anammox was sustainable throughout the model domain and all three functional types were able to coexist within the coexistence domain which contains the net zero N loss point. To determine which of the two model scenarios is more likely, observations could test to see whether anammox strains subsist in NO_2_^-^-limited environments (e.g. Bristow et al. 2016), and experiments could probe the competition of anammox (ideally, of low NO_2_^-^ adapted strains) versus heterotrophs for NO_2_^-^.

The coexistence of all three functional types (DNRA, anammox, and denitrification) has been observed in natural systems. For example, Kalvelage et al. (2013) and Roland et al. (2017) both measured simultaneous DNRA, anammox, and denitrification in coastal OMZs and Lake Kivu, respectively. The narrow range over which the model was able to simulate coexistence of the three functional types is likely due to the strict competitive exclusion of the simple model. This differs from the environment, where density-dependent mortality, due to ‘kill-the-winner’ grazing (Thingstad 2000) or viral lysis, may prevent complete competitive exclusion. Alternatively, anammox could, in principle, co-exist with the other metabolisms when neither OM or N are limiting, as has been demonstrated in instances of coupled and synergistic DNRA and anammox consortia (Han et al. 2020, Valiente et al. 2022, White et al. 2023, Zhou, Zhao, and Zhuang 2023).

The model presented here is simplified by design, aimed at broad relevance, and its complexity could be expanded in many ways, depending on the research question. For example, the complexity of OM substrates provides another axis over which to study the ecological outcomes of the model. Here, for simplicity, we have considered one bulk pool of OM with a particular stoichiometry that reflects the open ocean (C_6.6_H_10.9_O_2.6_N, Anderson 1995). Using a different or else simpler C source (such as glucose or ethanol) would alter the proportion of energy available for biomass synthesis, changing estimates of biomass yields. While the relative competitiveness of the included functional types in our model would not be altered significantly by OM stoichiometry, the numerical value of the OM:NO_2_^-^ supply ratios (such as those displayed along the x axis of Fig. 4) would change accordingly.

Our model is designed to indicate broad, regionally and globally relevant patterns (Giblin et al. 2013, Xu et al. 2025), and so we do not expect our model to be able to capture all of the heterogeneity of empirical observations. The current model does not resolve many of the complexities inherent in natural systems, such as niche heterogeneity, complex mortality dynamics (such as viral lysis, which would dampen competitive exclusion), chemoautotrophic and non-canonical DNRA and denitrification (Zhang et al. 2023, Wen et al. 2025), and metabolic flexibility which may vary the internal redox constraint for some species (Vuono et al. 2019). Our framework could be extended to consider some of these additional aspects, such as the chemoautotrophic analogues and mixotrophic denitrification (Zhang et al. 2015, Zhang et al. 2023). The model’s plasticity provides an adaptable tool for understanding the role that DNRA and denitrification competition plays in determining retention and loss of inorganic N across globally distributed and diverse anoxic water columns.

Because the microbially explicit model captures microbial feedbacks dynamically, it may be particularly helpful for understanding the ecosystem scale biogeochemical impacts of eutrophication. By better understanding the positive feedback loop in which eutrophication promotes DNRA and then DNRA, in turn, produces bioavailable NH_4_^+^, applied aquatic ecologists may be able to use evidence of a shift from denitrification to DNRA dominance as an indicator of impending trophic regime shift. This would allow for the identification of ecosystem change at the early stages of eutrophication before the proliferation of large and potentially ecologically deleterious algal blooms. Similarly, by mechanistically explaining these feedbacks, our results can improve our projections of N loss versus retention in aquatic systems in response to eutrophication and other anthropogenic perturbations.

## CONCLUSION

By basing our model on a fundamental predictor of microbial metabolism (the underlying redox chemistry), we provide a theoretically grounded and computationally rigorous understanding of anaerobic N-cycling microbial communities in anoxic water columns without dependency on site specific data or species-specific traits. This paves a way towards an improved prediction of globally relevant biogeochemical fluxes in unobserved environments (including future conditions). We identified the transition point at which inorganic N retention by DNRA balances inorganic N loss by denitrification (and also anammox), resulting in net zero N loss. The relative loss or retention of bioavailable N is of critical importance to N limited aquatic ecosystems, both inland and marine.

The model provides a universal understanding of the controls on N retention versus loss that also resolves functional microbial biomass. With additional efforts, the model outputs may be converted to and compared with microbial abundances inferred from sequencing data as well as measured per cell activity rates. Therefore, the presented model may contribute to extant global biogeochemical models as well as provide a tool for hypothesis generation and testing across scales.

## ACKNOWLEDGEMENTS

This work and the resulting manuscript have benefited from the feedback of the Zakem lab group at Carnegie Science. JMF was supported by the Simons Postdoctoral Fellowship in Marine Microbial Ecology (Award #LS-FMME-00993455), as was XS (Award #LS-FMME-00871981). EJZ acknowledges support from the National Science Foundation (Grant #2125142), the Simons Early Career Investigator in Aquatic Microbial Ecology and Evolution Award (Award #LS-ECIAMEE-00006631), and the European Research Council (ERC) Synergy Grant (RECLESS, #101167445)

## DATA AVAILABILITY

Code is available on GitHub at https://github.com/jfadum/DNRA-vs-Denit/tree/main

## Notes

### Competing Interest Statement

The authors have declared no competing interest.

## REFERENCES

Algar, Christopher K., and Joseph J. Vallino. 2014. “Predicting Microbial Nitrate Reduction Pathways in Coastal Sediments.” Aquatic Microbial Ecology 71 (3): 223–38. 10.3354/ame01678.

Anderson, Laurence A. 1995. “On the Hydrogen and Oxygen Content of Marine Phytoplankton.” Deep Sea Research Part I: Oceanographic Research Papers 42 (9): 1675–80. 10.1016/0967-0637(95)00072-E.

Anil, Akpinar, Charria Guillaume, Caroline Cusack, Fernandez Vicente, and Diarmuid O’Conchubhair. 2018. “Gap Analysis of Links between Coastal and Open Ocean Networks.” Report. AtlantOS. 10.13155/57443.

Babbin, Andrew R., Richard G. Keil, Allan H. Devol, and Bess B. Ward. 2014. “Organic Matter Stoichiometry, Flux, and Oxygen Control Nitrogen Loss in the Ocean.” Science 344 (6182): 406–8. 10.1126/science.1248364.

Bleakley, BH, and JM Tiedje. 1982. “Nitrous oxide production by organisms other than nitrifiers or denitrifiers.” Appl Environ Microbiol. 44(6):1342–8. doi: 10.1128/aem.44.6.1342-1348.1982.

Bristow, L., Callbeck, C., Larsen, M. et al. 2017. “N2 production rates limited by nitrite availability in the Bay of Bengal oxygen minimum zone.” Nature Geosci 10, 24–29. 10.1038/ngeo2847

Buchanan, Pearse J., Xin Sun, J. L. Weissman, Daniel McCoy, Daniele Bianchi, and Emily J. Zakem. 2025. “Oxygen Intrusions Sustain Aerobic Nitrite-Oxidizing Bacteria in Anoxic Marine Zones.” Science 388 (6751): 1069–74. 10.1126/science.ado0742.

Canfield, D.E. & Kraft, B. 2022. “The ‘oxygen’ in oxygen minimum zones.” Environmental Microbiology, 24(11), 5332–5344. 10.1111/1462-2920.16192

Cole, J. A., and C.M. Brown. 1980. “Nitrite Reduction to Ammonia by Fermentative Bacteria: A Short Circuit in the Biological Nitrogen Cycle.” FEMS Microbiology Letters 7 (2): 65–72. 10.1111/j.1574-6941.1980.tb01578.x.

Dalsgaard, T., Thamdrup, B., Farías, L., & Revsbech, N. P. 2012. “Anammox and denitrification in the oxygen minimum zone of the eastern South Pacific.” Limnology and Oceanography, 57(5), 1331–1346. 10.4319/lo.2012.57.5.1331

De Brabandere, Loreto, Don E. Canfield, Tage Dalsgaard, Gernot E. Friederich, Niels Peter Revsbech, Osvaldo Ulloa, and Bo Thamdrup. 2014. “Vertical Partitioning of Nitrogen-Loss Processes across the Oxic-Anoxic Interface of an Oceanic Oxygen Minimum Zone.” Environmental Microbiology 16 (10): 3041–54. 10.1111/1462-2920.12255.

Deininger, Anne, and Helene Frigstad. 2019. “Reevaluating the Role of Organic Matter Sources for Coastal Eutrophication, Oligotrophication, and Ecosystem Health.” Frontiers in Marine Science 6. https://www.frontiersin.org/articles/10.3389/fmars.2019.00210.

Diaz, Robert J. 2001. “Overview of Hypoxia around the World.” Journal of Environmental Quality 30 (2): 275–81. 10.2134/jeq2001.302275x.

Fadum, J. M., M. A. Borton, R. A. Daly, K. C. Wrighton, and E. K. Hall. 2024. “Dominant Nitrogen Metabolisms of a Warm, Seasonally Anoxic Freshwater Ecosystem Revealed Using Genome Resolved Metatranscriptomics.” mSystems 9 (2): e01059–23. 10.1128/msystems.01059-23.

Fadum, Jemma M., and Ed K. Hall. 2023. “Nitrogen Is Unlikely to Consistently Limit Primary Productivity in Most Tropical Lakes.” Ecosphere 14 (3): e4451. 10.1002/ecs2.4451.

Garcia-Robledo, Emilio, Cory C. Padilla, Montserrat Aldunate, Frank J. Stewart, Osvaldo Ulloa, Aurélien Paulmier, Gerald Gregori, and Niels Peter Revsbech. 2017. “Cryptic Oxygen Cycling in Anoxic Marine Zones.” Proceedings of the National Academy of Sciences 114 (31): 8319–24. 10.1073/pnas.1619844114.

Giblin, A.E., C.R. Tobias, B. Song, N. Weston, G.T. Banta, and V.H. Rivera-Monroy. 2013. “The Importance of Dissimilatory Nitrate Reduction to Ammonium (DNRA) in the Nitrogen Cycle of Coastal Ecosystems” 26 (3): 124–31. 10.5670/oceanog.2013.54.

Han, Xinkuan, Shuchan Peng, Lilan Zhang, Peili Lu, and Daijun Zhang. 2020. “The Co-Occurrence of DNRA and Anammox during the Anaerobic Degradation of Benzene under Denitrification.” Chemosphere 247 (May):125968. 10.1016/j.chemosphere.2020.125968.

IPCC, 2013: Climate Change 2013: The Physical Science Basis. Contribution of Working Group I to the Fifth Assessment Report of the Intergovernmental Panel on Climate Change [Stocker, T.F., D. Qin, G.-K. Plattner, M. Tignor, S.K. Allen, J. Boschung, A. Nauels, Y. Xia, V. Bex and P.M. Midgley (eds.)]. Cambridge University Press, Cambridge, United Kingdom and New York, NY, USA, 1535 pp.

Jia, Mingsheng, Mari K. H. Winkler, and Eveline I. P. Volcke. 2020. “Elucidating the Competition between Heterotrophic Denitrification and DNRA Using the Resource-Ratio Theory.” Environmental Science & Technology 54 (21): 13953–62. 10.1021/acs.est.0c01776.

Kalvelage, Tim, Gaute Lavik, Phyllis Lam, Sergio Contreras, Lionel Arteaga, Carolin R. Löscher, Andreas Oschlies, Aurélien Paulmier, Lothar Stramma, and Marcel M. M. Kuypers. 2013. “Nitrogen Cycling Driven by Organic Matter Export in the South Pacific Oxygen Minimum Zone.” Nature Geoscience 6 (3): 228–34. 10.1038/ngeo1739.

Kartal, Boran, Laura van Niftrik, Jan T. Keltjens, Huub J. M. Op den Camp, and Mike S. M. Jetten. 2012. “Anammox--Growth Physiology, Cell Biology, and Metabolism.” Advances in Microbial Physiology 60:211–62. 10.1016/B978-0-12-398264-3.00003-6.

Kiørboe, Thomas, André Visser, and Ken H Andersen. 2018. “A Trait-Based Approach to Ocean Ecology.” ICES Journal of Marine Science 75 (6): 1849–63. 10.1093/icesjms/fsy090.

Kleiner, Manuel, Erin Thorson, Christine E. Sharp, Xiaoli Dong, Dan Liu, Carmen Li, and Marc Strous. 2017. “Assessing Species Biomass Contributions in Microbial Communities via Metaproteomics.” Nature Communications 8 (1): 1558. 10.1038/s41467-017-01544-x.

Koeve, W., and P. Kähler. 2010. “Heterotrophic Denitrification vs. Autotrophic Anammox – Quantifying Collateral Effects on the Oceanic Carbon Cycle.” Biogeosciences 7 (8): 2327–37. 10.5194/bg-7-2327-2010.

Kraft, Beate, Marc Strous, and Halina E. Tegetmeyer. 2011. “Microbial Nitrate Respiration – Genes, Enzymes and Environmental Distribution.” Journal of Biotechnology, New Frontiers in Microbial Genome Research, 155 (1): 104–17. 10.1016/j.jbiotec.2010.12.025.

Kraft, Beate, Halina E. Tegetmeyer, Ritin Sharma, Martin G. Klotz, Timothy G. Ferdelman, Robert L. Hettich, Jeanine S. Geelhoed, and Marc Strous. 2014. “The Environmental Controls That Govern the End Product of Bacterial Nitrate Respiration.” Science 345 (6197): 676–79. 10.1126/science.1254070.

Lam, Phyllis, Gaute Lavik, Marlene M. Jensen, Jack van de Vossenberg, Markus Schmid, Dagmar Woebken, Dimitri Gutiérrez, Rudolf Amann, Mike S. M. Jetten, and Marcel M. M. Kuypers. 2009. “Revising the Nitrogen Cycle in the Peruvian Oxygen Minimum Zone.” Proceedings of the National Academy of Sciences 106 (12): 4752–57. 10.1073/pnas.0812444106.

Litchman, Elena, Christopher A. Klausmeier, Oscar M. Schofield, and Paul G. Falkowski. 2007. “The Role of Functional Traits and Trade-Offs in Structuring Phytoplankton Communities: Scaling from Cellular to Ecosystem Level.” Ecology Letters 10 (12): 1170–81. 10.1111/j.1461-0248.2007.01117.x.

Lotti, T., R. Kleerebezem, C. Lubello, and M. C. M. van Loosdrecht. 2014. “Physiological and Kinetic Characterization of a Suspended Cell Anammox Culture.” Water Research 60 (September):1–14. 10.1016/j.watres.2014.04.017.

Mccollom, T. M., and J. P. Amend. 2005. “A Thermodynamic Assessment of Energy Requirements for Biomass Synthesis by Chemolithoautotrophic Micro-Organisms in Oxic and Anoxic Environments.” Geobiology 3 (2): 135–44. 10.1111/j.1472-4669.2005.00045.x.

Meiler, S., Britten, G.L., Dutkiewicz, S., Gradoville, M.R., Moisander, P.H., Jahn, O. and Follows, M.J. 2022. “Constraining uncertainties of diazotroph biogeography from nifH gene abundance.” Limnol Oceanogr, 67: 816–829. 10.1002/lno.12036

Moore, C. M., M. M. Mills, K. R. Arrigo, I. Berman-Frank, L. Bopp, P. W. Boyd, E. D. Galbraith, et al. 2013. “Processes and Patterns of Oceanic Nutrient Limitation.” Nature Geoscience 6 (9): 701–10. 10.1038/ngeo1765.

Pajares, Silvia, Martín Merino-Ibarra, Miroslav Macek, and Javier Alcocer. 2017. “Vertical and Seasonal Distribution of Picoplankton and Functional Nitrogen Genes in a High-Altitude Warm-Monomictic Tropical Lake.” Freshwater Biology 62 (7): 1180–93. 10.1111/fwb.12935.

Rittman, Bruce, and Perry McCarty. 2001. Environmental Biotechnology: Principles and Applications. McGraw-Hill Education.

Roland, Fleur A. E., François Darchambeau, Alberto V. Borges, Cédric Morana, Loreto De Brabandere, Bo Thamdrup, and Sean A. Crowe. 2018. “Denitrification, Anaerobic Ammonium Oxidation, and Dissimilatory Nitrate Reduction to Ammonium in an East African Great Lake (Lake Kivu).” Limnology and Oceanography 63 (2): 687–701. 10.1002/lno.10660.

Rütting, T., Boeckx, P., Müller, C., and Klemedtsson, L. 2011. “Assessment of the importance of dissimilatory nitrate reduction to ammonium for the terrestrial nitrogen cycle.” Biogeosciences, 8, 1779–1791, 10.5194/bg-8-1779-2011

Sinsabaugh, Robert L., Stefano Manzoni, Daryl L. Moorhead, and Andreas Richter. 2013. “Carbon Use Efficiency of Microbial Communities: Stoichiometry, Methodology and Modelling.” Ecology Letters 16 (7): 930–39. 10.1111/ele.12113.

Smets, Wenke, Jonathan W. Leff, Mark A. Bradford, Rebecca L. McCulley, Sarah Lebeer, and Noah Fierer. 2016. “A Method for Simultaneous Measurement of Soil Bacterial Abundances and Community Composition via 16S rRNA Gene Sequencing.” Soil Biology and Biochemistry 96 (May):145–51. 10.1016/j.soilbio.2016.02.003.

Smits, Adrianne P., Ed K. Hall, Bridget R. Deemer, Facundo Scordo, Carolina C. Barbosa, Stephanie M. Carlson, Kaelin Cawley, et al. 2025. “Too Much and Not Enough Data: Challenges and Solutions for Generating Information in Freshwater Research and Monitoring.” Ecosphere 16 (3): e70205. 10.1002/ecs2.70205.

Strohm, Tobin O., Ben Griffin, Walter G. Zumft, and Bernhard Schink. 2007. “Growth Yields in Bacterial Denitrification and Nitrate Ammonification.” Applied and Environmental Microbiology 73 (5): 1420–24. 10.1128/AEM.02508-06.

Sun, Xin, Pearse J. Buchanan, Irene H. Zhang, Magdalena San Roman, Andrew R. Babbin, and Emily J. Zakem. 2024. “Ecological Dynamics Explain Modular Denitrification in the Ocean.” Proceedings of the National Academy of Sciences 121 (52): e2417421121. 10.1073/pnas.2417421121.

Thingstad, T. Frede. 2000. “Elements of a Theory for the Mechanisms Controlling Abundance, Diversity, and Biogeochemical Role of Lytic Bacterial Viruses in Aquatic Systems.” Limnology and Oceanography 45 (6): 1320–28. 10.4319/lo.2000.45.6.1320.

Tiedje, James M., Alan J. Sexstone, David D. Myrold, and Joseph A. Robinson. 1983. “Denitrification: Ecological Niches, Competition and Survival.” Antonie van Leeuwenhoek 48 (6): 569–83. 10.1007/BF00399542.

Tilman, David. 1982. Resource Competition and Community Structure. (MPB-17), Volume 17. Princeton University Press. 10.2307/j.ctvx5wb72.

Tracey, John C., Andrew R. Babbin, Elizabeth Wallace, Xin Sun, Katherine L. DuRussel, Claudia Frey, Donald E. Martocello III, Tyler Tamasi, Sergey Oleynik, and Bess B. Ward. 2023. “All about Nitrite: Exploring Nitrite Sources and Sinks in the Eastern Tropical North Pacific Oxygen Minimum Zone.” Biogeosciences 20 (12): 2499–2523. 10.5194/bg-20-2499-2023.

Valiente, N., F. Jirsa, T. Hein, W. Wanek, J. Prommer, P. Bonin, and J. J. Gómez-Alday. 2022. “The Role of Coupled DNRA-Anammox during Nitrate Removal in a Highly Saline Lake.” Science of The Total Environment 806 (February):150726. 10.1016/j.scitotenv.2021.150726.

Vallino, J. J., C. S. Hopkinson, and J. E. Hobbie. 1996. “Modeling Bacterial Utilization of Dissolved Organic Matter: Optimization Replaces Monod Growth Kinetics.” Limnology and Oceanography 41 (8): 1591–1609. 10.4319/lo.1996.41.8.1591.

van den Berg, Eveline M., Marissa Boleij, J. Gijs Kuenen, Robbert Kleerebezem, and Mark C. M. van Loosdrecht. 2016. “DNRA and Denitrification Coexist over a Broad Range of Acetate/N-NO3ȓ Ratios, in a Chemostat Enrichment Culture.” Frontiers in Microbiology 7 (November). 10.3389/fmicb.2016.01842.

van den Berg, Eveline M., Julius L. Rombouts, J. Gijs Kuenen, Robbert Kleerebezem, and Mark C. M. van Loosdrecht. 2017. “Role of Nitrite in the Competition between Denitrification and DNRA in a Chemostat Enrichment Culture.” AMB Express 7 (1): 91. 10.1186/s13568-017-0398-x.

Vuono, David C., Robert W. Read, James Hemp, Benjamin W. Sullivan, John A. Arnone, Iva Neveux, Robert R. Blank, et al. 2019. “Resource Concentration Modulates the Fate of Dissimilated Nitrogen in a Dual-Pathway Actinobacterium.” Frontiers in Microbiology 10 (January). 10.3389/fmicb.2019.00003.

Ward, B., Devol, A., Rich, J. et al. 2009. “Denitrification as the dominant nitrogen loss process in the Arabian Sea.” Nature 461, 78–81. 10.1038/nature08276

Wen, Wan-Ru, Tian-Chen Liu, Sheng-Qiang Fan, Xin Tan, Yang Lu, De-Feng Xing, Bing-Feng Liu, Nan-Qi Ren, and Guo-Jun Xie. 2025. “Dissimilatory Nitrate Reduction to Ammonium Driven by Different Electron Donors: Mechanisms, Recent Advances, and Future Perspectives.” Chemical Engineering Journal 507 (March):160625. 10.1016/j.cej.2025.160625.

White, Christian, Edmund Antell, Sarah L. Schwartz, Jennifer E. Lawrence, Ray Keren, Lijie Zhou, Ke Yu, Weiqin Zhuang, and Lisa Alvarez-Cohen. 2023. “Synergistic Interactions between Anammox and Dissimilatory Nitrate Reducing Bacteria Sustains Reactor Performance across Variable Nitrogen Loading Ratios.” Frontiers in Microbiology 14 (August):1243410. 10.3389/fmicb.2023.1243410.

Wright, Jody J., Kishori M. Konwar, and Steven J. Hallam. 2012. “Microbial Ecology of Expanding Oxygen Minimum Zones.” Nature Reviews Microbiology 10 (6): 381–94. 10.1038/nrmicro2778.

Xu, Xiaoguang, Yuxuan Yang, Yiwen Zhou, Jie Ma, Jining Li, Xiaohong Zhou, Xiaoli Zhao, Fengchang Wu, and Kang Song. 2025. “Global Patterns and Drivers of Coupling between Anammox and Denitrification Processes across Inland Aquatic Ecosystems.” Communications Earth & Environment 6 (1): 1–11. 10.1038/s43247-024-01980-w.

Zakem, Emily J., Amala Mahadevan, Jonathan M. Lauderdale, and Michael J. Follows. 2020. “Stable Aerobic and Anaerobic Coexistence in Anoxic Marine Zones.” The ISME Journal 14 (1): 288–301. 10.1038/s41396-019-0523-8.

Zakem, Emily J., Martin F. Polz, and Michael J. Follows. 2020. “Redox-Informed Models of Global Biogeochemical Cycles.” Nature Communications 11 (1): 5680. 10.1038/s41467-020-19454-w.

Zhang, Irene H., Xin Sun, Amal Jayakumar, Samantha G. Fortin, Bess B. Ward, and Andrew R. Babbin. 2023. “Partitioning of the Denitrification Pathway and Other Nitrite Metabolisms within Global Oxygen Deficient Zones.” ISME Communications 3 (1): 1–14. 10.1038/s43705-023-00284-y.

Zhang, Jingxu, Yuyin Yang, Lei Zhao, Yuzhao Li, Shuguang Xie, and Yong Liu. 2015. “Distribution of Sediment Bacterial and Archaeal Communities in Plateau Freshwater Lakes.” Applied Microbiology and Biotechnology 99 (7): 3291–3302. 10.1007/s00253-014-6262-x.

Zhou, Lijie, Bikai Zhao, and Wei-Qin Zhuang. 2023. “Double-Edged Sword Effects of Dissimilatory Nitrate Reduction to Ammonium (DNRA) Bacteria on Anammox Bacteria Performance in an MBR Reactor.” Water Research 233 (April):119754. 10.1016/j.watres.2023.119754.

