## Supporting Information: Figures for "Redox-constrained microbial ecology dictates nitrogen loss versus retention"

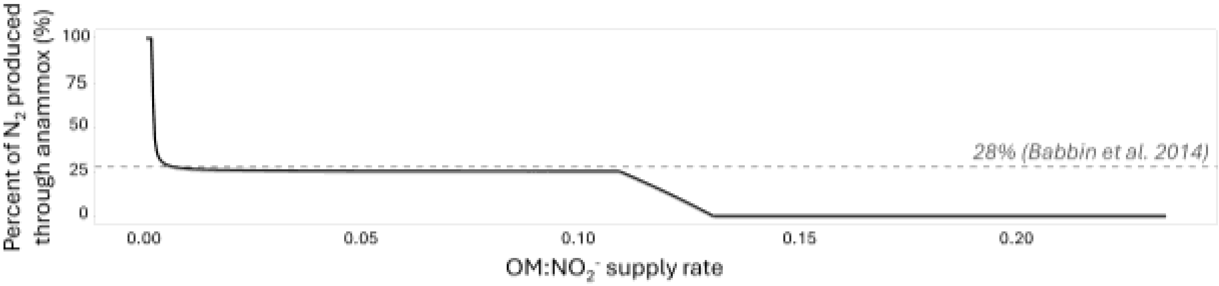


**Figure S1.** Percentage of $N_2$ generated in the chemostat by anammox (from both the reduction of $NO_2^-$ and oxidation of $NH_4^+$).

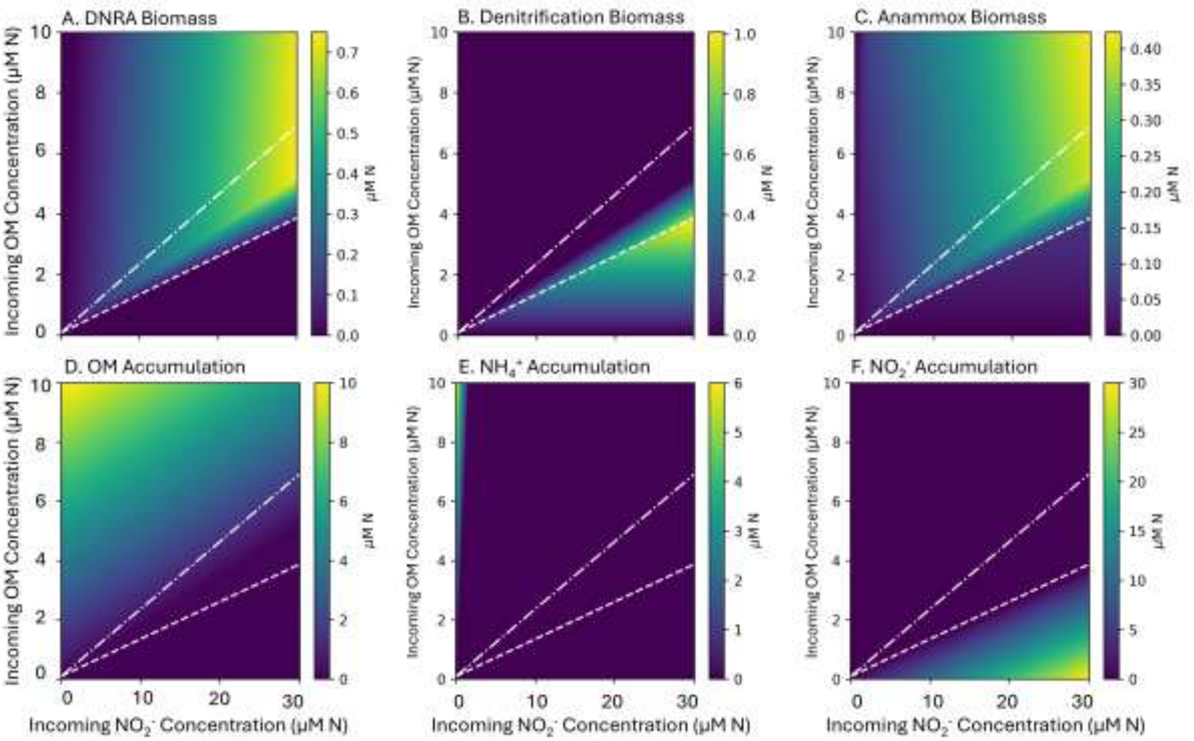


**Figure S2.** *Increased competitiveness of anammox scenario:* Multiple steady-state solutions of the ecosystem model when anammox has the lowest $NO_2$* with anaerobic N-cycling metabolic functional types across varying supply of OM and $NO_2^-$, showing biomass of A) DNRA, B) Denitrification ($NO_2^- \rightarrow N_2O + N_2O \rightarrow N_2$) and C) Anammox. Chemostat accumulation (steady-state concentrations) of D) OM, E) $NH_4^+$ with F) $NO_2^-$. The white lines indicate the consumption vectors for DNRA (− • − • −) and denitrification (---). Axis supply rates indicate resource supplies without dilution (0.04).

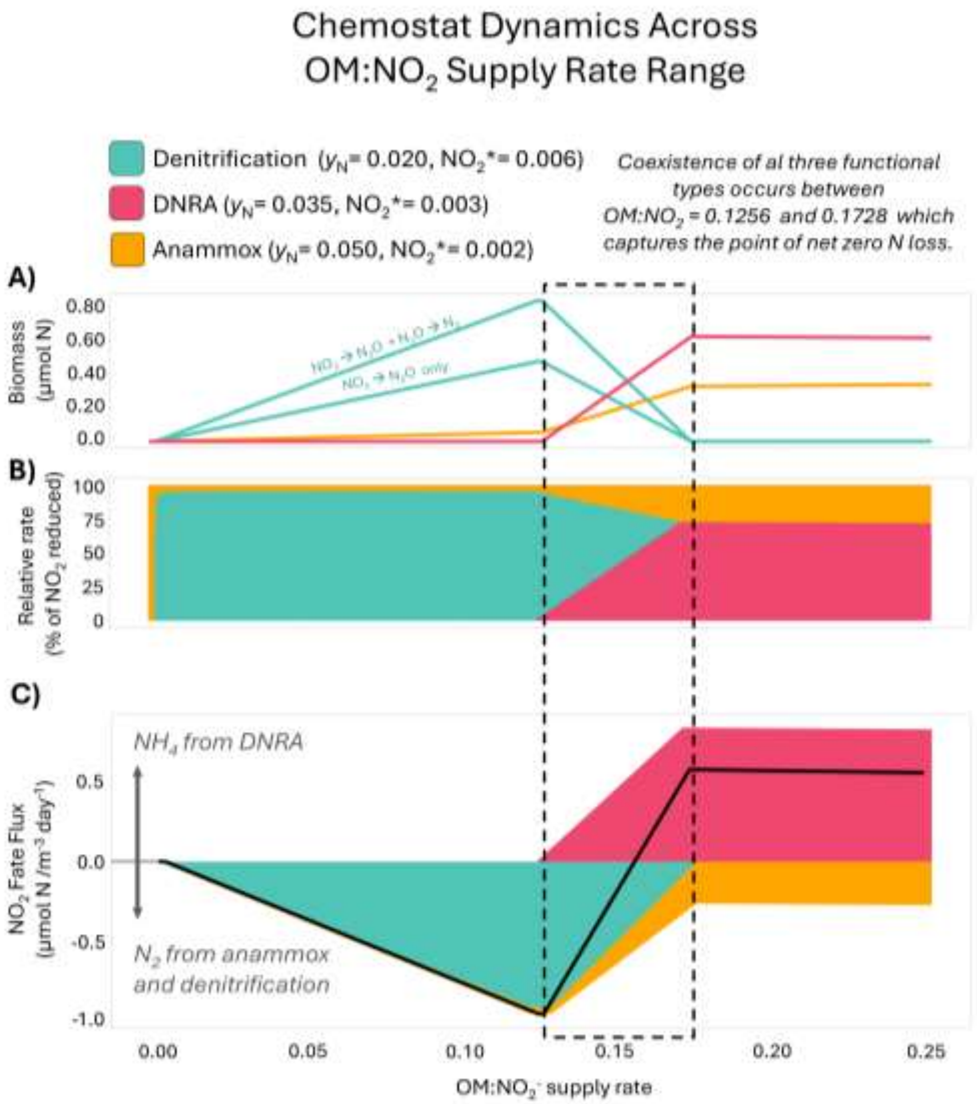


**Figure S3.** *Increased competitiveness of anammox scenario:* Ecological and biogeochemical outcomes governing the fate of $NO_2^-$ in the ecosystem model when anammox has the lowest $NO_2$* as a function of the relative supply of OM to $NO_2^-$ (here specifically with incoming $NO_2^-$ concentration of 25 µM N and variable OM supply rate). A) Biomasses of DNRA, denitrification, and anammox functional types, B) Percentage of $NO_2^-$ reduced via the three functional types, C) $N_2$ and $NH_4^+$ production with solid black line denoting net balance (different than default model because balance includes N loss from anammox). Range of coexistence for all three functional types contained in dashed line box.

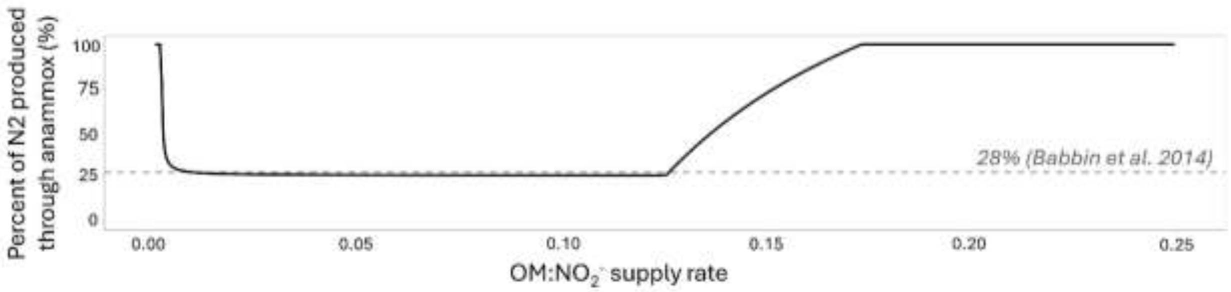


**Figure S4.** *Increased competitiveness of anammox scenario:* Percentage of $N_2$ generated in the chemostat by anammox (from both the reduction of $NO_2^-$ and oxidation of $NH_4^+$) when anammox has the lowest $NO_2$*.

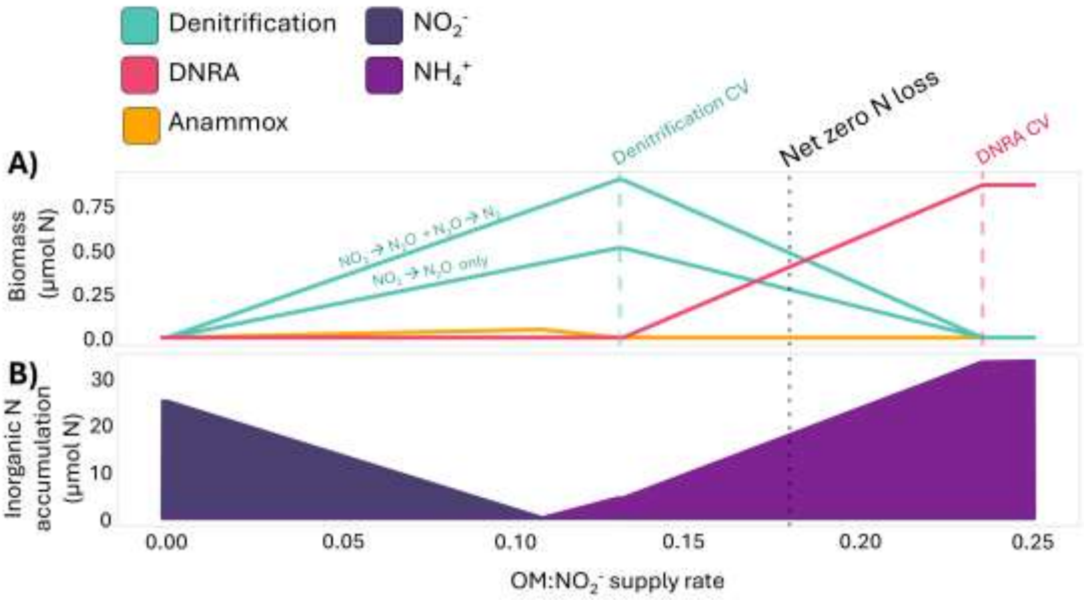


**Figure S5.** Ecological and biogeochemical outcomes governing the fate of $NO_2^-$ in the ecosystem model (default version) as a function of the relative supply of OM to $NO_2^-$ (here specifically with incoming $NO_2^-$ concentration of 25 µM N and OM supply rate varying). A) Biomasses of DNRA, denitrification, and anammox functional types, same as Figure 4A in main text for reference, B) Speciation of inorganic N accumulation.
