## Supporting Information: Tables for "Redox-constrained microbial ecology dictates nitrogen loss versus retention"

**Table S1.** Fixed values in model

| | |
|---|---|
| OM incoming concentration range | 0-10 µM N, increments of 0.01 µM N |
| $NO_2^-$ incoming concentration range | 0-30 µM N, increments of 0.1 µM N (Fig. 3); 25 µM N (Fig. 4) |
| Dilution rate | 0.04 /day (Sun et al., 2024) |
| Simulation total time | 2000 days (ensuring equilibrium has been reached) |
| Timestep for numerical integration | 0.001 day |

**Table S2.** Key kinetic parameters for metabolic functional types.

| Substrate | Parameter | Value | Source(s) |
|---|---|---|---|
| OM | Maximum uptake rate ($V^{max}$)<br>Half-saturation conc. ($K_{OM}$) | 3.0 mols of organic N/ mol N biomass/day<br>1.0 µM | (Jia et al., 2020; Zakem et al., 2020) |
| $NO_2$ | Maximum uptake rate ($V^{max}$)<br>Half-saturation conc. ($K_N$) | 30 mols of N/ mol N biomass/day<br>0.1 µM | (Awata et al., 2013; Sun et al., 2024) |
| $N_2O$ | Maximum uptake rate ($V^{max}$)<br>Half-saturation conc. ($K_{N2O}$) | 30 mols of N/ mol N biomass/day<br>0.6 µM | (Sun et al., 2017) |

**References**


Anderson, L. A. (1995). On the hydrogen and oxygen content of marine phytoplankton. *Deep Sea Research Part I: Oceanographic Research Papers*, *42*(9), 1675–1680. https://doi.org/10.1016/0967-0637(95)00072-E

Awata, T., Oshiki, M., Kindaichi, T., Ozaki, N., Ohashi, A., & Okabe, S. (2013). Physiological Characterization of an Anaerobic Ammonium-Oxidizing Bacterium Belonging to the "Candidatus Scalindua" Group. *Applied and Environmental Microbiology*, *79*(13), 4145–4148. https://doi.org/10.1128/AEM.00056-13

Jia, M., Winkler, M. K. H., & Volcke, E. I. P. (2020). Elucidating the Competition between Heterotrophic Denitrification and DNRA Using the Resource-Ratio Theory. *Environmental Science & Technology*, *54*(21), 13953–13962. https://doi.org/10.1021/acs.est.0c01776

Rittman, B., & McCarty, P. (2001). *Environmental Biotechnology: Principles and Applications*. McGraw-Hill Education.

Sun, X., Buchanan, P. J., Zhang, I. H., San Roman, M., Babbin, A. R., & Zakem, E. J. (2024). Ecological dynamics explain modular denitrification in the ocean. *Proceedings of the National Academy of Sciences*, *121*(52), e2417421121. https://doi.org/10.1073/pnas.2417421121

Zakem, E. J., Mahadevan, A., Lauderdale, J. M., & Follows, M. J. (2020). Stable aerobic and anaerobic coexistence in anoxic marine zones. *The ISME Journal*, *14*(1), Article 1. https://doi.org/10.1038/s41396-019-0523-8
