## Supporting Information: Text for "Redox-constrained microbial ecology dictates nitrogen loss versus retention"

### Text S1: DNRA functional type

Following $C_{c_{OM}}H_{h_{OM}}O_{o_{OM}}N_{n_{OM}}$ and $C_{c_B}H_{h_B}O_{o_B}N_{n_B}$ to describe the composition of organic matter (OM) and biomass (B), respectively, the denominator *d* normalizes all reactions to one electron (Rittman & McCarty 2001).

$$d_{OM} = 4c_{OM} + h_{OM} - 2o_{OM} - 3n_{OM}$$

$$d_B = 4c_B + h_B - 2o_B - 3n_B$$

Ignoring $H_2O$ and $H^+$, the full reaction for DNRA is:

$$\frac{1}{d_{OM}}OM + \frac{1-f}{6}NO_2^- \rightarrow \frac{f}{d_B}B + \left(\frac{c_{OM}}{d_{OM}} - \frac{c_B \cdot f}{d_B}\right)CO_2 + \left(\frac{n_{OM}}{d_{OM}} - \frac{n_B \cdot f}{d_B} + \frac{1-f}{6}\right)NH_4^+$$

where *f* is the fraction of the electrons from the electron donor (OM) used in biomass synthesis (thus 1-*f* describes electron allocation to respiration). For full methods for determining *f* from the redox reactions of N-cycling heterotrophs, see Sun et al., 2024. Briefly, the *f* for DNRA is estimated by balancing the free energy gained from the redox reaction with an empirical estimate of the free energy required for biomass synthesis, constrained by the observed range of baseline growth efficiencies for heterotrophs as mentioned in the main text (0.2—0.3 mol B/mol OM). The resulting value of *f* for DNRA (0.10) is ~60% of that for denitrification ($NO_2^- \rightarrow N_2O$). For a full description of the denitrification and anammox types, see Sun et al., 2024.

### Text S2: Calculating $NH_4^+$ supply from implied $NO_3^-$ reduction

Anammox requires $NH_4^+$ as its electron donor. Without supplying $NH_4^+$ to the chemostat, anammox would be entirely dependent on the $NH_4^+$ produced from DNRA and the oxidation of OM by denitrification. Therefore, in order to have a more realistic representation of anammox function, we supply the $NH_4^+$ from the implied $NO_3^-$ reduction.

Following the metabolic budget for $NO_3^-$ reduction from Sun et al. 2024, $NH_4^+$ is produced in direct proportion to OM utilization in $NO_3^-$ reduction to $NO_2^-$ as:

$$\frac{1}{d_{OM}}OM + \frac{1}{2}NO_3^- \rightarrow \frac{1}{d_{OM}}NH_4^+ + \frac{1}{2}NO_2^-$$

When both $NO_3^-$ and $NO_2^-$ reducing functional types (Sun et al. 2024) coexist in the OM-limited regime (where anammox is competitive in the default model), 60% of the OM is oxidized by the $NO_3^-$ reducer. Therefore, we supply $NH_4^+$ to the chemostat at 60% of the OM supply rate.

### Text S3: Defining loss and retention for 'net zero N loss'

We define N retention as the $NH_4^+$ produced by DNRA from the reduction of $NO_x$, with growth rate, μ, and dilution rate, *D*:

$$\frac{1}{y_{N_{DNRA}}}\mu B_{DNRA} D$$

Other sources of $NH_4^+$production in the chemostat include a comparatively small amount of $NH_4^+$ produced by DNRA from its oxidation of OM, and similarly, the $NH_4^+$ produced by the oxidation of OM by denitrification. However, we are interested in the fate of $NO_2^-$ and therefore focus exclusively on the $NH_4^+$ produced through the reduction of $NO_2^-$.

Similarly, we define N loss from denitrification as the $N_2$ it produces, as:

$$\frac{1}{y_{N_{Denit}}}\mu B_{Denit} D$$

As $D$ is equal for both fluxes, when anammox is negligible (as in the default model), retention and loss balance at:

$$\frac{1}{y_{N_{Denit}}}\mu B_{Denit} = \frac{1}{y_{n_{DNRA}}}\mu B_{DNRA}$$ (provided as Eqn. 2 in main text)

### References


Rittman, Bruce, and Perry McCarty. 2001. *Environmental Biotechnology: Principles and Applications*. McGraw-Hill Education.

Sun, X., Buchanan, P. J., Zhang, I. H., San Roman, M., Babbin, A. R., & Zakem, E. J. (2024). Ecological dynamics explain modular denitrification in the ocean. *Proceedings of the National Academy of Sciences*, *121*(52), e2417421121. https://doi.org/10.1073/pnas.2417421121

Zakem, E. J., Mahadevan, A., Lauderdale, J. M., & Follows, M. J. (2020). Stable aerobic and anaerobic coexistence in anoxic marine zones. *The ISME Journal*, *14*(1), Article 1. https://doi.org/10.1038/s41396-019-0523-8
